# Active role of free RNA during the early phase of proteostasis stress

**DOI:** 10.1101/564799

**Authors:** Marion Alriquet, Adrían Martínez-Limón, Gerd Hanspach, Martin Hengesbach, Gian G. Tartaglia, Giulia Calloni, R. Martin Vabulas

## Abstract

Transient sequestration of proteins and RNA is an essential principle of cellular reaction to stress. Compared to polypeptides, less is known about the role of RNA released from polysomes during acute proteostasis stress. Using quantitative mass spectrometry, we identified a set of proteins assembled by free RNA in the heat-shocked mammalian cytosol. RNA-associated proteins displayed higher disorder and larger size, which supports the role of multivalent interactions during the initial phase of the RNA granule formation. Structural features of the free RNA interactors defined them as a subset of RNA-binding proteins. The interactome contained preferentially the active form of eIF2α. The interaction between assembled proteins *in vivo* required RNA. The reconstitution of the association process *in vitro* indicated to the multimolecular basis for the increased binding to RNA upon heat shock in the cytosol. Our results reveal how free RNA can participate in reorganization of cellular functions during proteostasis stress.

## INTRODUCTION

Protein unfolding and aggregation during proteostasis stress can lead to the irreparable damage of the cellular proteome and cell death. One of the strategies eukaryotic cells use to survive under stress is the sequestration of proteins and RNA in transient membrane-less assemblies. Cytosolic stress granules (SGs) is the best known example of those structures. Low complexity (LC) of the amino acid sequence is characteristic of a fraction of proteins assembling into membrane-less structures ^1^. Another feature of the sequestering proteins is the multivalency of their interactions ^2^. Compared to proteins, less molecular details are known regarding the role of RNA during stress granulation. A recent study uncovered the RNA-seeded formation of nuclear amyloid bodies upon heat shock and during acidosis ^3^. It was reported that the secondary structure of mRNA affects the composition of membrane-less granules ^4^. Importantly, RNA binding can modulate phase separation and toxicity of solid-like assemblies ^5^.

The polypeptide-coding mRNAs are coated by specialized proteins at any stage of their lifecycle. More than 1000 proteins can associate with mRNAs to participate in processing, transport, translation and turnover of transcripts ^6^. Since many mRNA-binding proteins (RBPs) have LC domains ^7^, mRNA can effectively increase the local concentration of RBPs, thus facilitating the associated phase transition. Accordingly, a significant fraction of RBPs are found in mRNA-containing granules formed *in vivo* and *in vitro* ^8,9^. Although mRNAs lose the associated proteins during ribosomal translation, a dense packing and particular arrangement of ribosomes seem to reduce the exposure of stripped RNA to the cytosolic content ^10^. One of few circumstances when free mRNA appears in the cytosol is the disassembly of polysomes during stress-induced shutdown of protein synthesis. It has been proposed that the massive increase of free mRNA creates a relative insufficiency of RNA-stabilizing proteins such as YB-1 ^11^. Under these conditions, mRNAs would be able to associate with granulation-prone proteins, an initial step of SG formation. Here, we aimed at a molecular understanding of this early phase of the stress response. Quantitative mass spectrometry was used to identify proteins in heat shocked HeLa lysates which interact with free RNA baits. Usually, UV crosslinking is applied to stabilize direct protein-RNA interactions. We decided to avoid crosslinking and use softer washing procedures in order to reveal the complexity of the assemblies. Secondly, we used two defined RNA sequences, not the mix of all possible polyA-containing RNAs as it is done during standard interactomics studies. An impressive interactivity potential of a given RNA sequence was revealed. On the other hand, the significant overlap of the two interactomes argues for the common molecular mechanism of free RNA processing during stress.

## EXPERIMENTAL SECTION

### Reagents, Plasmids, Antibodies

All chemicals were from Sigma-Aldrich (Saint Louis, MO) unless otherwise indicated.

Flag-tagged TRMT6 and TRMT61A mammalian expression vectors were purchased from Genscript (Piscataway, NJ). Two additional Flag tags where inserted into the vector coding TRMT61A. Human HSP70 under T7 promoter was cloned into pCH16 vector (a kind gift of H.-C. Chang) for *in vitro* transcription. Human B-Raf under T7 promoter was a gift from Dustin Maly (Addgene #40775) and was further modified by the insertion of two additional FLAG tags and of an RBS sequence. The following antibodies were used: anti-TRMT6 (A303-008A-M) from Bethyl (Montgomery, TX); anti-TRMT61A (sc-107105) and anti-S6 (sc-74459) from Santa Cruz Biotechnology (Dallas, TX); anti-eIF2α (9722), anti-eIF2α (phosphor-Ser51) (3398), anti-eIF4E (9742), HRP-conjugated anti-rabbit-IgG (7074) and Alexa Fluor 647-conjugated anti-rabbit IgG (4414) from Cell Signaling (Danvers, MA); anti-Flag (F1804) and HRP-conjugated anti-mouse IgG (A9044) from Sigma-Aldrich.

### Cell culture

The human HeLa cell line was cultured in DMEM supplemented with 10% (vol/vol) FBS, 2 mM L-glutamine, 100 IU/mL penicillin G, 100 μg/mL streptomycin sulfate, and nonessential amino acids.

### RNA Electroporation and Imaging

4×10^6^ HeLa cells were transfected with 4 μg Cy5-RNA or the molar equivalent of Cy5-55mer oligo by electroporation in 400 µL intracellular buffer (135 mM KCl, 0,2 mM CaCl2, 2 mM MgCl_2_, 5 mM EGTA, 10 mM Hepes, pH 7.5) freshly supplemented with 2 mM ATP. Electroporation was perfomed at 310 V and 950 μF using a Gene Pulser Xcell system from Bio-Rad Laboratories (Hercules, CA). After electroporation, the cells were washed and plated in a 12-well plate with polylysine-coated cover slides at 1 million cells/well. They were left for recovery at 37°C for 6 h. Then cells were washed in PBS and fixed with 4% paraformaldehyde/PBS at RT for 10 min. One slide was then treated with 250 units Benzonase in PBS supplemented with 2 mM MgCl_2_ at 37°C for 20 min. All the slides were washed with PBS, stained with DAPI, mounted in PBS and imaged using a Zeiss LSM-780 inverted confocal microscope with a 63x oil immersion objective. The quantitative comparison was performed using CellProfiler ^12^ as follows. Nuclei were detected and cells were identified by propagation from the nuclei. Intensity of Cy5 fluorescence was measured for each cell and Cy5-positive cells were identified. The experiment was performed in triplicates and at least 300 cells were analyzed per condition and per repetition to determine the percentage of Cy5-positive cells.

### Analysis of Protein Translation Activity

DMEM medium without methionine and cysteine was supplemented with 25 mM HEPES NaOH pH 7.5, 100 µg/mL streptomycin sulphate, 2 mM L-glutamine, 0.2 mM cystine and 0.02 mM methionine and used to prepare a suspension of HeLa cells at 1×10^7^ cells/mL. Cells were pre-incubated for indicated time at 37°C or 45°C before 50 μCi/mL of [35S]-methionine (Hartmann Analytics, Germany) was added for 3 min labelling of nascent proteins. Samples were mixed immediately with hot SDS sample buffer, nucleic acids were sheared using a 26 G needle with at least ten strokes and run in 10% SDS-PAGE. Afterwards, the gel was dried at 70°C for 1 h and exposed on a phosphorimager plate overnight. Typhoon 9400 imaging system (GE Healthcare) was used to visualize and quantify the radioactive signal.

### Pulldowns, Immunoprecipitation, Western Blotting

6×10^7^ HeLa cells were resuspended in 2 mL medium and heat-shocked at 45°C for 1 h. After two PBS washes, the cells were resuspended in 1,5 volume of Hypotonic lysis buffer (HLB: 10 mM Hepes KOH pH 7.6, 10 mM K acetate, 1.5 mM Mg acetate, 2 mM DTT) and kept on ice for 10 min. The lysate was passed through a 20 G needle 20 times and centrifuged at 640 g for 5 min at 4°C. The supernatant was collected, supplemented with 1/10 volume of 1 M potassium acetate and centrifuged at 10.400 g for 20 min at 4°C. Protein concentration was normalized to 5 μg/μL. For each sample, 20 μL of Dynabeads M-280 Streptavidin (Thermo Fisher Scientific, Waltham, MA) were washed once with BW (5 mM Tris HCl pH 7.5, 0.5 mM EDTA, 1 M NaCl), twice with Solution A (0.1 M NaOH, 0.05 M NaCl), once with Solution B (0.1 M NaCl) and once with BW. The beads were loaded with 2 μg biotinylated HSP70 mRNA (or DEPC-treated H_2_O) in BW in a volume of 40 μL for 30 min at RT, washed three times with BW and once with HLB. 100 μL of lysate was added to the beads and incubated at 4°C for 2 h with gentle shaking. The beads were washed six times with 50 μL Washing Buffer (WB: 25 mM Tris HCl pH 7.5, 10 mM MgCl_2_, 2 mM DTT) and three times with 50 mM Tris HCl pH 7.4, 150 mM NaCl. The beads were frozen at −80°C until further processing for mass spectrometry. For biochemical verification, 100 μL of lysate were added to the beads and incubated at 4°C for 3 h with gentle shaking, then washed six times with 50 μL WB. Elution was performed with 250 units of Benzonase in 10 μL WB at 37°C for 15 min.

For co-immunoprecipitation, 2×10^6^ HeLa cells were transfected with 5 μg 3xFlag-TRMT61A and 5 μg Flag-TRMT6 or 10 μg pcDNA3.1-Hygro. The cells were collected 24 h later, resuspended in HEPES/DMEM and incubated at 37°C or 45°C for 1 h. Lysates without or with Benzonase treatment were prepared as described above, centrifuged at 10.400 g for 20 min at 4°C. Protein concentration in supernatants was normalized to 1,5 μg/μL. 10 μL anti-Flag M2 affinity gel were blocked with 1% BSA in HLB for 1 h on ice, incubated with 200 μL lysate at 4°C for 3 h, washed four times with WB and eluted with 50 μL 3xFLAG peptide solution at 100 μg/mL in TBS for 5 min at RT. Reducing SDS sample buffer was added to the eluates, samples were resolved using 10% SDS-PAGE and transferred onto nitrocellulose membranes. Membranes were blocked with 5% skim milk or 5% BSA in TBST (TBS/0.1% Tween 20), probed with the indicated antibodies and developed using the SuperSignal West Pico PLUS. Chemiluminescence images were acquired with the Chemidoc MP imaging system and bands quantified using the Image Lab 5.0 software.

### PolyT Pulldowns

5×10^6^ HeLa cells were resuspended in 1 mL DMEM supplemented with 25 mM Hepes NaOH pH 7.5 and incubated at 45°C for 60 min, while control cells were kept at 37°C. The cells were washed and lysed in 300 µL lysis buffer (LB: 10 mM Tris HCl pH 7.4, 100 mM KCl, 5 mM MgCl_2,_ 0.5 % sodium deoxycholate, 1% Triton X-100) supplemented with 2 mM DTT, Phosphatase Inhibitor Cocktail 2 and RNAsin for 5 min at RT. Lysates were passed 5 times through a 20 G needle, protein concentration was measured and normalized. 400 µL of lysate at the protein concentration of 1,5 µg/µL were added to 100 μL of washed Oligo d(T)25 Magnetic Beads (NEB, Ipswich, MA) and incubated at RT for 30 min. 2/3 of the beads were washed 3x with LB and resuspended in 10 µL LB supplemented with 250 units of Benzonase nuclease for protein elution. After 15 min at 37°C, the supernatant was collected and analyzed by SDS-PAGE and western blotting using anti-TRMT6 antibody. The rest of the beads were processed for RNA elution. For this, the beads were washed 2x with WB-I (20 mM Tris HCl pH7.5, 500 mM LiCl, 0.1% LiDS, 1 mM EDTA, 5 mM DTT), 2x with WB-II (20 mM Tris-HCl pH 7.5, 500 mM LiCl, 1 mM EDTA) and 1x with Low Salt Buffer (20 mM Tris HCl pH 7.5, 200 mM LiCl, 1 mM EDTA). The beads were eluted with 30 μL of 20 mM Tris HCl pH 7.5, 1 mM EDTA at 50°C for 2 min. RNA amounts were determined with RiboGreen and used for normalization.

### Fluorescence Microscopy

HeLa cells were seeded in a 12-well plate at 10^5^ cells/well on a polylysine-coated cover slides. Next day, medium was refreshed and supplemented with 25 mM Hepes. Cells were heat-shocked at 45°C for 1 h or treated with 1 mM arsenite for 30 min at 37°C. Cells were washed with PBS, fixed with 3.7% paraformaldehyde, permeabilized with acetone at −20°C for 5 min, blocked with 1% BSA/PBS for 1 h at RT and incubated with anti-eIF4E for 1 h. Incubation with anti-Rabbit-IgG Alexa Fluor647 conjugate for 1 h, three washes with PBS and staining with DAPI followed. Slides were imaged using a Zeiss LSM-780 inverted confocal microscope with a 63x oil immersion objective. Quantification was performed using CellProfiler. First, nuclei were detected, then cells were identified by propagation from the nuclei. Stress granules were identified and counted. The experiment was performed in triplicates and at least 100 cells were analyzed per condition and per repetition.

### Polysome and 80S Monosome Analyses

HeLa cells were seeded at 5×10^6^ per 10 cm-dish. Next day, DMEM was refreshed and supplemented with 25 mM Hepes. The dishes were incubated at 45°C for 1 h or treated with 1 mM arsenite for 30 min at 37°C. The cells were washed twice with ice-cold PBS supplemented with 100 μg/mL cycloheximide and resuspended in 600 μL lysis buffer (10 mM Tris HCl pH 7.4, 100 mM KCl, 5 mM MgCl2, 0,5% sodium deoxycholate, 1% Triton X-100) supplemented with Phosphatase inhibitor cocktail 2 (1:100), RNasin (1:1000) and 100 μg/mL cycloheximide. After a 15 min incubation on ice, the lysates were centrifuged at 10.000 g for 5 min at 4°C. 550 μL of supernatant was loaded on a cold 10-50% Sucrose gradient in an Open-top Polyclear 16,8 mL tube (Seton, Egelsbach, Germany) and centrifuged in an Optima XPN-80 ultracentrifuge form Beckman Coulter (Brea, CA) using a SW28.1 rotor, at 100.000 g for 4 h at 4°C. Gradient absorbance at 254 nm was measured on a Gradient station (Biocomp) at a speed of 0.2 mm/second.

For monosome analysis, the 80S fraction was collected. Samples were normalized according to the absorbance at 260 nm and analyzed by western blotting using anti-S6 antibody.

### RNA Synthesis and Biotinylation *In Vitro*

For *in vitro* transcription, linearized plasmids (HSP70 using Xho I, BRaf using Xba I) were purified by phenol/chloroform/isoamylalcohol extraction and ethanol precipitation. Ampliscribe T7 high Yield kit (Epicenter, Madison, WI) was used to transcribe 1 μg of template for 4 h at 37°C. The reaction mix was digested with DNaseI and RNA was purified. If the RNA had to be further biotinylated, it was purified by phenol/chloroform/isoamylalcohol followed by ethanol precipitation and resuspended in a small volume of DEPC-treated water. Otherwise, RNA was purified using MegaClear transcription clean-up kit (Thermo Fisher Scientific). Quality of the mRNA was assessed on a formaldehyde agarose gel. 50 pmol of *in vitro* transcribed RNA was biotinylated using RNA 3’ End biotinylation Kit (Thermo Fisher Scientific) overnight at 16°C. Biotinylated RNA was purified using MegaClear kit.

### 5’-labelling of RNA

For Cy5-labeling of RNA, GMP was incorporated during *in vitro* transcription by adding it to the transcription reaction in excess at 15 mM. The reaction was incubated at 37°C for 4 h, digested with DNaseI and purified using Qiagen RNeasy Mini kit. Quality of the mRNA was assessed by running an aliquote on a Formaldehyde agarose gel. <u>EDC coupling</u>: The RNA was dissolved in 75 μL 1x Reaction Buffer (150 mM NaCl, 10 mM EDTA, 10 mM Potassium phosphate, pH 7) and added to a tube containing 12.5 mg EDC (1-Ethyl-3-(3-dimethylaminopropyl)carbodiimid*hydrochloride). 50 μL of 0.25 M ethylene diamine, 0.1 M imidazole in water were immediately added to the reaction. The tube was vortexed and briefly centrifuged, then 200 μL of 0.1 M imidazole (pH 6) solution were added. The reaction incubated at 37°C for 2.5-3 h. The RNA was ethanol precipitated twice in 0.3 M NaOAc (pH 5.5). <u>Dye coupling:</u> One Cy5 Mono-Reactive Dye Pack was used for up to three labeling reactions. The dye was dissolved in 60 μL RNase-free DMSO and split in 20 μL aliquots. The RNA was dissolved in 20 μL of a freshly prepared 0.1 M NaHCO_3_ solution. One 20 μL aliquot was added to the RNA and the mixture was incubated for 90 min at RT in the dark. The RNA was ethanol precipitated twice in 0.3 M NaOAc (pH 5.5), resuspended in water and stored at −20 °C.

### tRNA Synthesis *In Vitro*

Human initiation tRNA (iMet) was synthesized *in vitro* using Ampliscribe T7 High Yield transcription kit. Two oligonucleotides were used at 1 µM for this purpose. One oligonucleotide (3’-T-TCG-AAT-TAT-GCT-GAG-TGA-TAT-<u>C</u>CG-TCT-CAC-CGC-GTC-GCC-TTC-GCA-CGA-CCC-GGG-TAT-TGG-GTC-TCC-AGC-TAC-CTA-GCT-TTG-GTA-GGA-GAC-G<u>G</u>T-GGT-5’) was used as a template. The resulting tRNA starts with G instead of A. The complementary base in the acceptor stem was accordingly exchanged into C. Both changes in the template are underlined. The second oligonucleotide (5’-A-AGC-TTA-ATA-CGA-CTC-ACT-ATA-3’) was used to prime the T7 RNA polymerase. The reaction was carried at 37°C for 4.5 h. The final product was purified by Phenol/Chloroform/Isoamylalcohol extraction and Ethanol precipitation.

### Protein Purification

Human his-tagged TRMT6/61A dimer was purified from Escherichia coli BL21 cells using 1 mL HisTrap HP affinity chromatography columns (GE Healthcare). Eluted fractions containing the recombinant proteins were pooled and incubated overnight with PreScission protease at 4°C. Protein solution was then subjected to size-exclusion chromatography using a HiLoad Superdex 200 column (GE Healthcare) in 50 mM HEPES NaOH pH 7.5, 150 mM NaCl and 1 mM DTT.

### Protein Melting Analysis

A fluorescent dye-based thermal melting assay was carried out using a CFX96 Real-Time PCR detection system. 10 μM of purified recombinant heterodimer TRMT6/61A were mixed with or without 10 μM tRNA and with SYPRO Orange dye (1:5000) in 100 mM Tris HCl pH 7.6, 100 mM NH_4_Cl, 10 mM MgCl_2_, 0.1 mM EDTA, and 1 mM DTT (added fresh). Fluorescence change was recorded while gradually increasing the temperature from 20°C to 80°C in 0.5°C/15 s steps (excitation at 450-490 nm, detection at 510-530 nm). Background fluorescence from the buffer as well as from the tRNA were subtracted before further analysis. The apparent melting temperature values were obtained by calculating the first derivative of the fluorescence change.

### CD Spectroscopy

CD spectra of proteins were recorded using a Jasco J-810 spectropolarimeter at 37°C and 45°C in 50 mM HEPES NaOH pH 7.5 and 150 mM NaCl. Proteins at 5 μM were placed in a 1-mm path cell (Hellma Analytics, Müllheim, Germany) and samples were scanned three times from 260 nm to 200 nm in 0.5 nm steps at 50 nm/min. The values from the buffer were subtracted.

### Protease Sensitivity Analysis

Recombinant proteins were diluted to 1.5 μM in 25 mM Hepes KOH pH 7.5, 150 mM NaCl, 5 mM CaCl_2_ and 2 mM DTT and incubated for 20 min at the indicated temperature. Proteinase K was added at 0,1 ng/μL and samples were collected after 3, 6 and 9 min. Hydrolysis was stopped by addition of PMSF followed by hot SDS sample buffer. The samples were analyzed by SDS-PAGE and Coomassie BB G-250 (CBB) staining. Images were acquired with the Chemidoc MP imaging system and bands quantified using the Image Lab 5.0 software (both from Bio-Rad).

### TRMT6/61A Binding to RNA *In Vitro*

*In vitro* transcribed tRNA was heated at 85°C for 3 min and then refolded in 100 mM Tris HCl pH 7.6, 100 mM NH_4_Cl, 10 mM MgCl_2_, 0.1 mM EDTA and 1 mM DTT at 30°C for 15 min. Recombinant TRMT6/61A was diluted to 10 nM in hypotonic lysis buffer supplemented with RNasin and phosphatase inhibitors and incubated at indicated temperature for 20 min. tRNA was added at a final concentration of 1 µM and the mixture was further incubated at room temperature for 20 min with 20 μL M-280 Streptavidin beads coated with biotinylated RNA. The beads were washed and eluted as described for RNA pulldowns.

### TRMT6/61A Binding to tRNA *In Vitro*

M200 Pro Microplate Reader (Tecan, Männedorf, Switzerland) was used to record the change in tRNA UV light absorbance upon binding to the TRMT6/61A heterodimer. tRNA (1 µM) and protein solution was prepared in 100 mM Tris HCl pH 7.6, 100 mM NH_4_Cl, 10 mM MgCl_2_, 0.1 mM EDTA, and 1 mM DTT (added fresh). The mix was pre-incubated for 10 min at 25°C in a UV-transparent 96-well plates (Corning). Then temperature was raised to 37°C or 42°C for another 10 min before measuring the absorbance between 230 nm and 350 nm in 2-nm steps. The formula (λmax - λ260)/λmax was used to determine the binding of the tRNA to the protein, where λmax is the 260 nm absorbance of the pure tRNA and λ260 is the 260 nm absorbance from the tRNA protein mix. The background absorbance from the buffer as well as from the protein only solution was subtracted before calculations.

### Quantitative Mass Spectrometry

#### Sample preparation

Pulled down proteins were processed on-beads for LC-MS/MS analysis as following. Beads were re-suspended in 50 µL 8M urea/50 mM Tris HCl pH 8.5, reduced with 10 mM DTT for 30 min and alkylated with 40 mM chloroacetamide for 20 min at 22°C. Urea was diluted to a final concentration of 2 M with 25 mM Tris HCl pH 8.5, 10% acetonitrile and proteins were digested with trypsin/lys-C mix overnight at 22°C. Acidified peptides (0.1% trifluoroacetic acid) were desalted and fractionated on combined C18/SCX stage tips (3 fractions). Peptides were dried and resolved in 1% acetonitrile, 0.1% formic acid.

#### LC-MS/MS

LC-MS/MS was performed on a Q Exactive Plus equipped with an ultra-high pressure liquid chromatography unit (Easy-nLC1000) and a Nanospray Flex Ion-Source (all three from Thermo Fisher Scientific). Peptides were separated on an in-house packed column (100 μm inner diameter, 30 cm length, 2.4 μm Reprosil-Pur C18 resin) using a gradient from mobile phase A (4% acetonitrile, 0.1% formic acid) to 30% mobile phase B (80% acetonitrile, 0.1% formic acid) for 60 min followed by a second step to 60% B for 30 min, with a flow rate of 300 nl/min. MS data were recorded in data-dependent mode selecting the 10 most abundant precursor ions for HCD with a normalized collision energy of 30. The full MS scan range was set from 300 to 2000 m/z with a resolution of 70000. Ions with a charge ≥2 were selected for MS/MS scan with a resolution of 17500 and an isolation window of 2 m/z. The maximum ion injection time for the survey scan and the MS/MS scans was 120 ms, and the ion target values were set to 3×10^6^ and 10^5^, respectively. Dynamic exclusion of selected ions was set to 30 s. Data were acquired using Xcalibur software.

#### Data analysis with MaxQuant

MS raw files from five (Hsp70 coding region interactome) or three biological replicates (B-Raf coding region interactome) of pulldown and background samples were analyzed with Max Quant (version 1.5.3.30) (Cox and Mann, 2008) using default parameters. Enzyme specificity was set to trypsin and lysC and a maximum of 2 missed cleavages were allowed. A minimal peptide length of 7 amino acids was required. Carbamidomethylcysteine was set as a fixed modification, while N-terminal acetylation and methionine oxidation were set as variable modifications. The spectra were searched against the UniProtKB human FASTA database (downloaded in November 2015, 70075 entries) for protein identification with a false discovery rate of 1%. Unidentified features were matched between runs in a time window of 2 min. In the case of identified peptides that were shared between two or more proteins, these were combined and reported in protein group. Hits in three categories (false positives, only identified by site, and known contaminants) were excluded from further analysis. For label-free quantification (LFQ), the minimum ratio count was set to 1.

#### Data analysis with Perseus

Bioinformatic data analysis was performed using Perseus (version 1.5.2.6) (Tyanova et al., 2016). Proteins identified in the pulldown experiments were further included in the analysis if they were quantified in at least 4 out of 5 biological replicates in at least one group (pulldown/background) for the Hsp70 coding region RNA and in at least 3 out of 3 biological replicates in at least one group (pulldown/background) for the B-Raf coding region RNA control. Missing LFQ values were imputed on the basis of normal distribution with a width of 0.3 and a downshift of 1.8. Proteins enriched in the pulldown (RNA over background binding) were identified by two-sample t-test at a permutation-based FDR cutoff of 0.001 and s0 = 0.1 for the Hsp70 coding region interactome and at a p-value cutoff of 0.01 for the B-Raf coding region interactome. Categorical annotations were added in Perseus and a Fisher’s exact test with a p-value threshold of 0.001 was run for GO term enrichment analysis.

### Bioinformatics Analysis

pI and molecular weight values were calculated using the pI/MW tool on the ExPASy website. Hydrophobicity and disorder propensity were analyzed with the boxplotter function of cleverSuite (Klus et al., 2014) using Kyte and Doolittle (Kyte and Doolittle, 1982) and TOP-IDB (Campen et al., 2008) scales, respectively. To analyze aggregation propensity the Zyggregator algorithm was used (Tartaglia and Vendruscolo, 2010). To analyze granulation propensity the catGRANULE algorithm was used (Bolognesi et al., 2016). Hsp70 mRNA interactors were compared to the human proteome as provided by Perseus (20592 proteins) and to the human RBPs (Gerstberger et al., 2014). Statistical significance of observed differences was assessed by Mann-Whitney test.

### Statistical Analysis

All repetitions in this study were independent biological repetitions performed at least three times if not specified differently. To identify significantly increased proteins in pulldowns (RNA over background binding) in mass spectrometry analyses, a two-sample t-test analysis of grouped biological replicates was performed using a FDR cutoff of 0.001 with s0 = 0.1 and a p-value cutoff of 0.01 for the Hsp70 and B-Raf coding region interactome, respectively. Means and standard deviations were calculated from at least three independent experiments.

## RESULTS

### Free RNA Forms Granules *In Vivo*

The majority of mRNA-specific regulatory associations are anchored on the untranslated regions (UTRs) of transcripts. Specifically, all known regulation of the HSP70 mRNA operates through these regions ^13^. To reduce the mRNA-specific bias, UTRs were deleted from the mRNAs used in this study (Figure 1A). Electroporation of *in vitro* transcribed and 5’-labelled RNA led to the efficient granule formation in up to 95% of transfected human HeLa cells (Figure 1B). The granulation reassured us in our strategy to reduce sequence specificity of the RNA, since native HSP70 mRNA is known to escape granulation ^14^. The formed structures contained RNA because nuclease treatment before imaging significantly reduced the percentage of cells containing them (Figure S1A). Cy5-labeled oligonucleotide was used to exclude that the fluorophore freed from RNA by intracellular hydrolysis accumulates to fluorescent foci in the cytosol. (Figure S1A).

**Figure 1.**
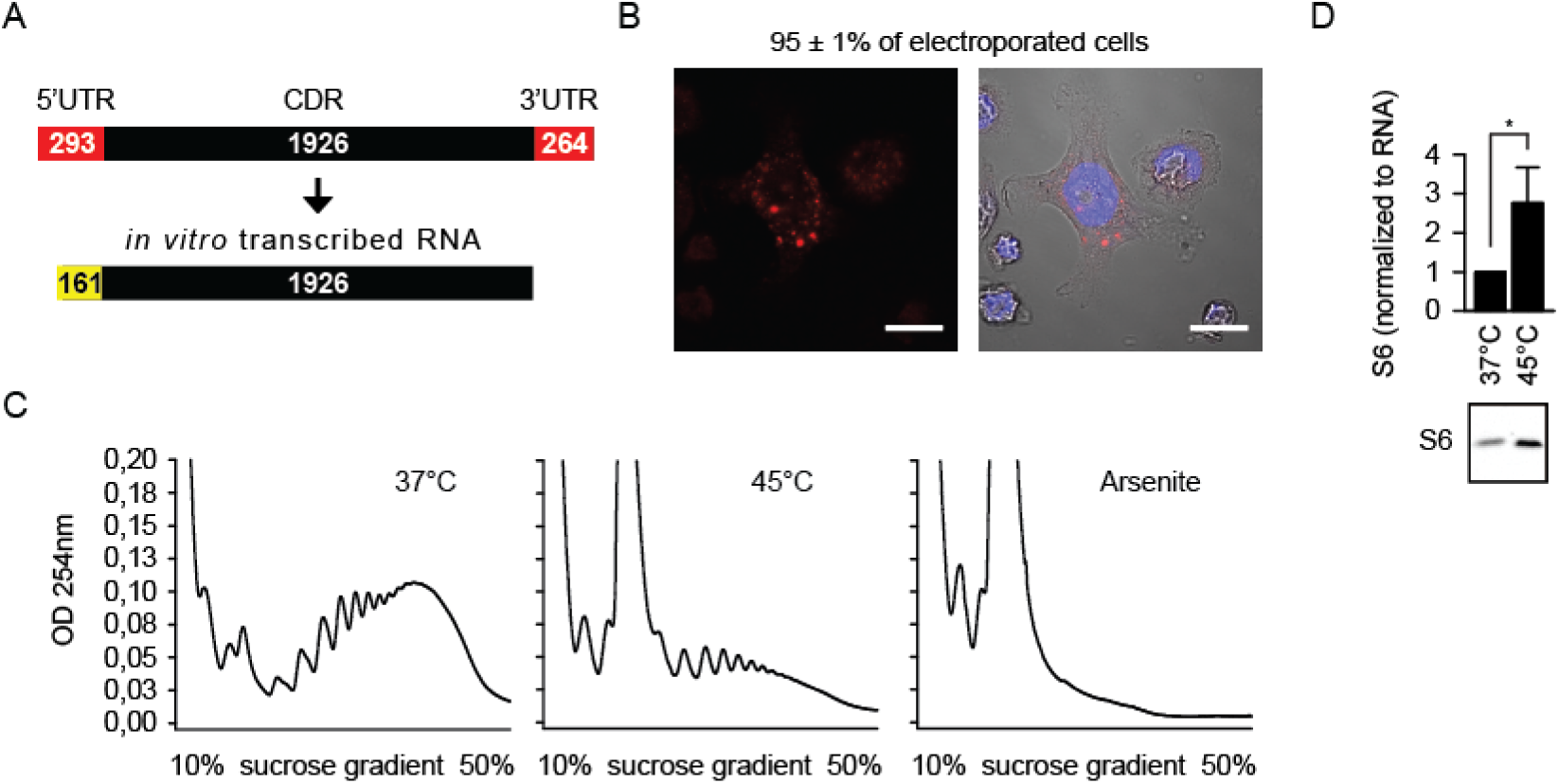
Free RNA Forms Granules *In Vivo*. (A) Schematic depiction of the RNA (gene *HSPA1A*) used in the study. The nucleotide length of different elements is indicated. UTR, untranslated region; CDR, coding region. Yellow, an additional fragment from the expression vector. (B) HeLa cells 6 h after electroporation with Cy5 (red) 5’-labelled RNA. DAPI staining (blue), nuclei. Scale bar 20 μm. Fraction of transfected cells with granules is indicated (N=3 independent experiments). Right, merged image. (C) Polysome profile under the indicated conditions. One representative out of three independent experiments is shown. (D) Ribosome amount in the monosome fraction as measured by anti-S6 western blot. *p<0.05, two-tailed t-test; N=3 independent experiments (mean + SD).

We then used the *in vitro* transcribed and biotin-labelled mRNA as a bait to identify RNA interactors in the heat-shocked HeLa cytosol, under conditions of polysome disassembly, free RNA release and stress granule (SG) formation. Cell treatment with 1 mM arsenite was used as a reference. The polysome profile indicated an early stage of cellular reorganization (Figure 1C), because it was reduced, yet not fully absent as in the case of the arsenite treatment. The monosome peak (rRNAs plus associated mRNAs) was strongly increased (Figure 1C) but had higher fraction of ribosomes (higher rRNA fraction means smaller fraction of mRNA per absorbance unit) as measured by anti-S6 western blot (Figure 1D). This observation confirmed that mRNAs from disassembled polysomes did not remain associated with ribosomes but were freed into the cytosol. Microscopically, imminent formation of stress granules (SG) was observed confirming the early phase of stress response (Figure S1B).

### Free RNA Assembles a Set of Proteins in the Heat-Shocked Cytosol

The bait RNA was incubated with the lysate of heat shocked cells and associated proteins were identified by means of mass spectrometry (Figure 2A and Table S1). We did not perform experiments with non-shocked lysates because they are not supposed to have free RNAs. Instead, we used binding to uncoated beads as background control. Five independent biological repetitions yielded highly reproducible label-free quantification of RNA interacting proteins (Figure S2A). When averaged and corrected for background binding to beads without RNA, a distinct set of 79 interacting proteins unveiled as visualized by the volcano plot (Figure 2B). More than half of the interactors (46) had been shown or are predicted to be RNA-binders ^15^ (Table S2). At the same time, proteostasis and signaling networks were represented by a number of proteins, such as the cochaperone DNAJC21, ubiquitin ligase RBB6, APC protein or casein kinase I.

**Figure 2.**
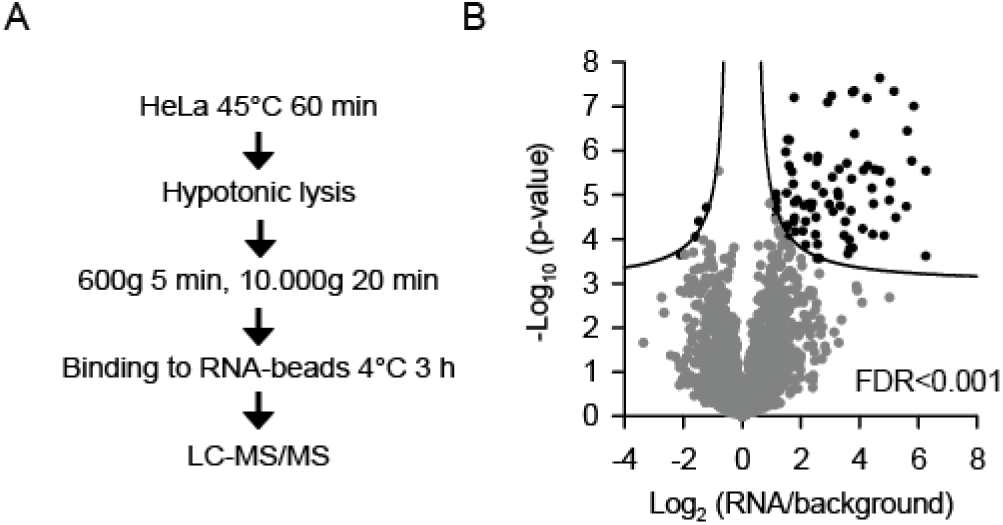
Free RNA Assembles a Set of Proteins in the Heat-Shocked Cytosol. (A) Experimental steps to identify the free RNA-assembled machinery in the heat shocked mammalian cytosol. (B) Volcano plot of quantified proteins plotted according to their enrichment on RNA over background with the statistical significance of the respective ratios plotted on the y-axis. False discovery rate (FDR) used to define the 79 interactors (black symbols) is indicated. N=5 independent experiments.

To reassure that the interactome was not dominated by the sequence identity, we determined the interactome of another UTR-deleted RNA of similar length (2295 nucleotides of the human BRaf coding region plus 184 nucleotides from the vector). This independent RNA is further called ‘control RNA’ (Table S3). With three repetitions the analysis was less deep, nevertheless, we identified 58 free RNA-interacting proteins in the heat-shocked cells (Figure S2B and Table S4). Importantly, ca. half of the interactome of the control RNA overlapped with the interactome of the first bait RNA with the high significance of p<4.7 × 10^−63^ based on hypergeometric distribution (Figure S2C).

### Interactors with Free RNA Display Distinct Structural Features

A possible concern while analyzing nucleic acid interactors are the trivial electrostatic interactions between negatively charged RNA and positively charged proteins. The analysis of the isoelectric point (pI) distribution showed only a slight increase of the mean pI value of the interactors above the proteome average (Figure 3A). This finding suggests that the electrostatics did not critically steer the associations. In contrast, the molecular size distribution was strongly shifted when compared to the distribution of the whole human proteome (Figure 3B). An average interactor is 74 kDa big while the human proteins have a molecular weight of 38 kDa on average (Figure 3C). The fractional difference was especially obvious for proteins larger than 100 kDa. The large size is indicative of a multidomain structure of the respective proteins and thus suggests the importance of multivalent interactions in the identified assembly.

**Figure 3.**
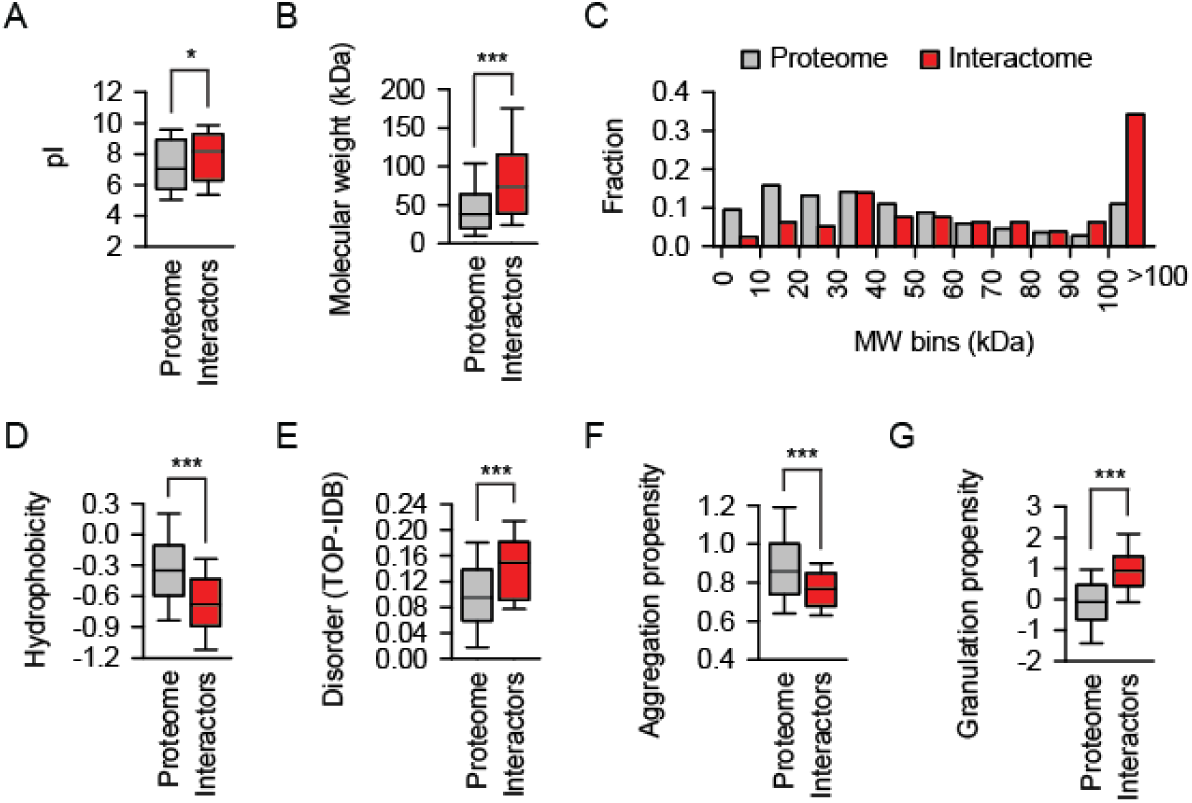
Interactors with Free RNA Display Distinct Structural Features. Distribution of isoelectric points (A), molecular weights (B and C), Kyte-Doolittle hydrophobicity scores (D), predicted disorder (E), aggregation propensity (F) and granulation propensity (G) scores of the indicated sets of proteins. *p<0.05, ***p<0.001, Mann-Whitney test.

The mean hydrophobicity calculated according to the Kyte-Doolittle scale was significantly lower for the RNA interactors (Figure 3D). A low hydrophobicity and lack of a hydrophobic core usually correlate with the lack of a defined structure. Consequently, a significant increase of structural disorder was predicted for the RNA interactors (Figure 3E). These properties might partially explain the reduced aggregation propensity of the interactors (Figure 3F), because, the presence of a hydrophobic core is a key requirement for aggregation. Finally, we used the recently developed algorithm *cat*GRANULE which estimates the granule-forming propensity of proteins ^16^. This parameter also turned out to be strongly discriminatory between the human proteome and the RNA interactors (Figure 3G).

Gene ontology (GO) analysis revealed an expected highly significant enrichment of nucleic acid/RNA binding categories among the 79 interactors (Figure S3A). Two further categories, “nucleolus” and “non-membrane-bound organelle”, pointed to the involvement of the interactome in the membrane-less protein sequestration, presumably in its early phase since the lysates were prepared from cells with few nascent SGs (Figure S1B). The comparison of the human RNA binding proteins ^15^ with the identified free-RNA interactome of heat shocked cells revealed differences between these groups at several structural parameters, including the granulation propensity (Figures S3B-S3G).

### Heat Shock Promotes RNA-Protein Association

Attenuation of the cap-dependent protein translation is an obligatory reaction to different stress stimuli, including the conformational stress that causes protein misfolding. eIF2α phosphorylation on serine 51 (S51) is the key to the shutdown of translation and the assembly of SG ^17^. We were intrigued to discover all three subunits of the eIF2 among the RNA interactors in the heat shocked lysate (Figure 4A). The enrichment of the individual proteins over the background was not high, yet significant, and came along with extensive sequence coverage by mass spectrometry (Figure S4A). The interaction could be verified biochemically (Figure 4B) and was found to increase significantly during heat shock. The association cannot be considered trivial, because the current translation initiation model states that eIF2 binds mRNAs after a complex with the 40S ribosome and eIF3 is formed. Neither 40S nor eIF3 were found in the interactome.

**Figure 4.**
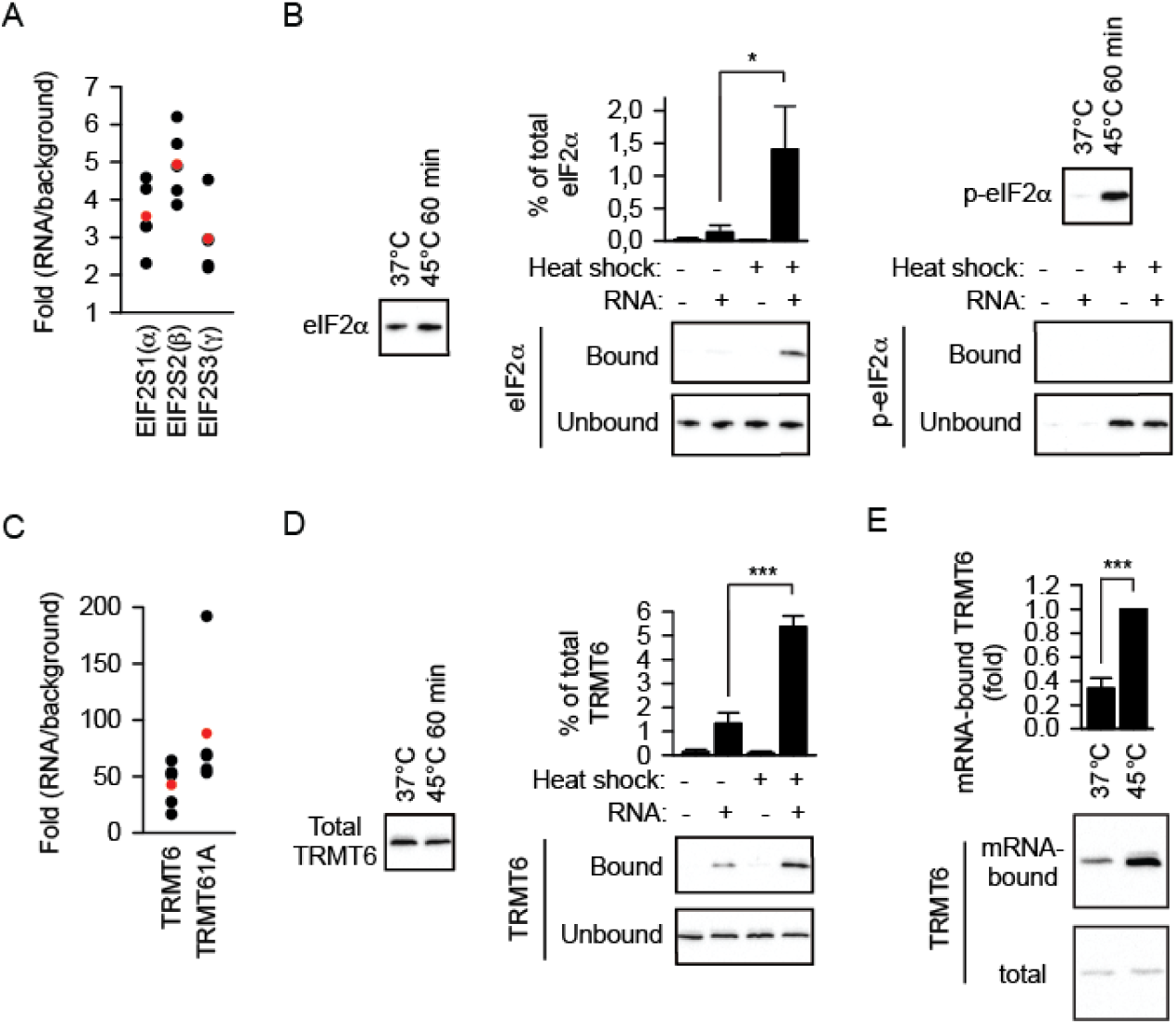
Heat Shock Promotes RNA-Protein Association in HeLa cells. (A) Enrichment values of the subunits of eIF2 in individual experiments (black symbols) and the enrichment averages (red symbols). (B) Left panels, amounts of total eIF2α and its serine 51-phosporylated form (p-eIF2α) as determined by western blotting. Right panels, enrichment of non-phosphorylated eIF2α on RNA upon heat shock. *p<0.05, two-tailed t-test; N=3 independent experiments (mean + SD). (C) Enrichment values of the subunits of the m^1^A methyltransferase TRMT6/61A in individual experiments (black symbols) and the enrichment averages (red symbols). (D) Upper panel, amounts of TRMT6 as determined by western blotting. Lower panel, enrichment of TRMT6 on RNA upon heat shock. ***p<0.001, two-tailed t-test; N=3 independent experiments (mean + SD). (E) Increased association of TRMT6 with mRNAs during heat shock as determined by western blot analysis of PolyT pulldowns. ***p<0.001, two-tailed t-test, N=3 independent experiments (mean + SD).

Notably, only the S51-non phosphorylated eIF2α could be detected in the pulldowns. The phospho-S51 eIF2α, through a tight interaction with eIF2B, leads to slower GDP-to-GTP exchange which results in an impaired translation initiation. Control experiments showed that the phosphorylated eIF2α accumulated in the lysate as expected (Figure S4B) along with an efficient shutdown of the bulk protein translation (Figure S4C). Thus, association of the active fraction of eIF2 on free RNA in the heat shocked lysates suggested a functional relevance of the interaction.

Next, we analyzed the TRMT6/61A complex, one of the top enrichments among the RNA interactors (Table S2, Figure 4C and Figure S4D). TRMT6/61A has been known as a tRNA methyltransferase able to methylate adenine 58 at N1 in tRNAs ^18^. Recently, m^1^A was identified also in eukaryotic mRNAs which implicated the modification in posttranscriptional regulation of gene expression ^19,20^. Two studies established TRMT6/61A as an m^1^A writer on mRNAs ^21,22^. Similarly to eIF2, heat shock enhanced the docking of TRMT6/61A on free mRNA (Figure 4D). TRMT6 increasingly associated with endogenous mRNA during heat shock (Figure 4E).

The heat shock increased the association of both complexes either directly or via common binding partners (Figure 5A). Hydrolysis of RNA in the lysate abolished this association (Figure 5B) which suggested that free RNA provides a scaffold to assemble regulatory components of the translation machinery during the early phase of the stress response.

**Figure 5.**
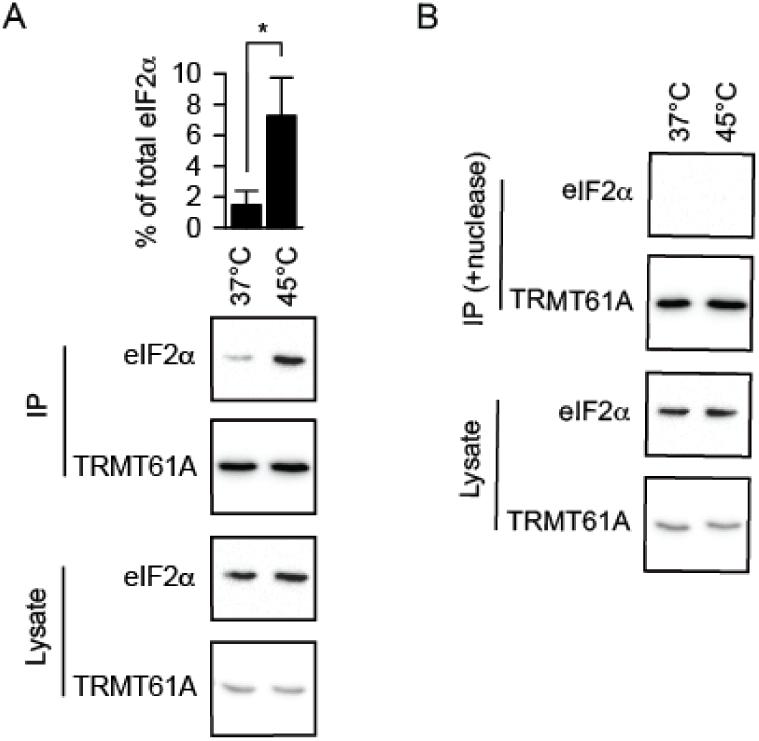
RNA Scaffolds Protein Complexes in Heat-Shocked HeLa cells. (A) Increased association of eIF2α with TRMT61A-containing complexes during heat shock as determined by immunoprecipitation (IP) of Flag-tagged TRMT6/61A. *p<0.05, two-tailed t-test; N=3 independent experiments (mean + SD). (B) RNA hydrolysis (+nuclease) abolishes association of eIF2α with the TRMT61A-containing complexes. One representative out of three independent experiments is shown.

Importantly, subunits of the eIF2 and TRMT6/61A were identified as significant interactors of the control RNA as well (Table S4).

### Structural Changes of TRMT6/61A at Higher Temperature

To understand the reasons for the heat-activated association of TRMT6/61A with the free RNA, the recombinant methyltransferase was purified (Figure S5A) and analyzed *in vitro*. The stability analysis of the proteins TRMT6/61A revealed two melting points (Figure 6A). The major unfolding happened around 51°C (T_m_). However, at temperature around 43°C, an additional partial transition could be detected (T_m’_). The presence of a TRMT6/61A substrate, the unmethylated tRNA, slightly increased T_m_ but decreased T_m’_ (Figure 6A, right panel). The early transition did not involve a substantial loss of secondary structure (Figure S5B). The lack of an appreciable difference in the circular dichroism (CD) spectrum suggested two possibilities. From one side, the T_m’_ at ca. 43°C might reflect the complex dissociation. TRMT6/61A exists as a tetramer formed by two heterodimers ^23^. tRNA binds across the dimer interface and involves TRMT6 and TRMT61A from the opposing heterodimers. tRNA was able to shift the major T_m_ (Figure 6A) which indicates intactness of the tRNA binding site up to at least 51°C and argues against the tetramer dissociation into dimers at lower temperature. The interface and predicted free energy between the monomers in the heterodimer are almost two times larger than those between the heterodimers in the tetramer ^23^. This makes the dissociation of heterodimers into monomers at 43°C even less probable. Alternatively, a subtle local unfolding might take place at this temperature. Analysis of the protease sensitivity of the complex supported the latter interpretation (Figure 6B). Interestingly, only the subunit TRMT6 showed an increased sensitivity.

**Figure 6.**
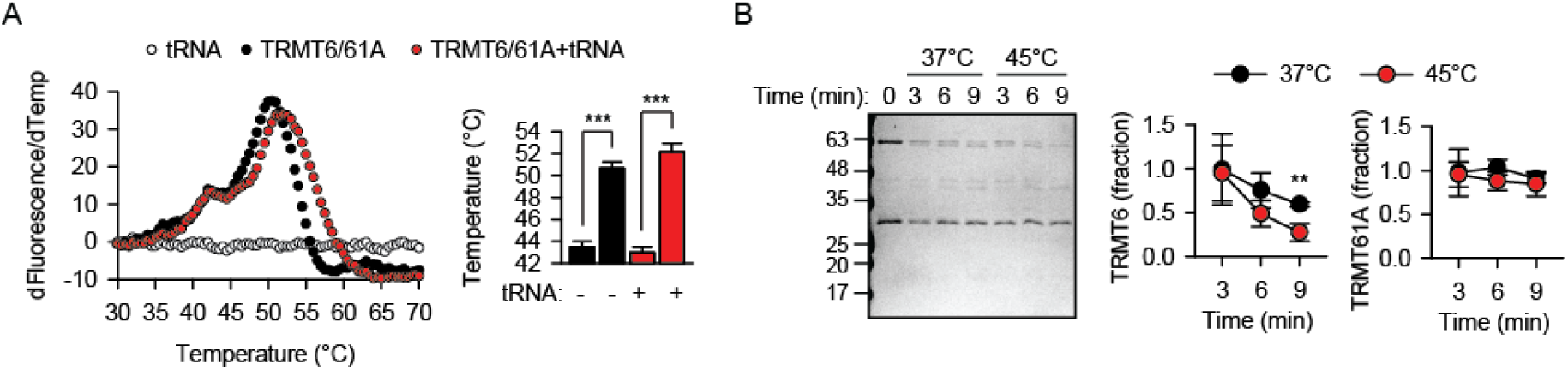
TRMT6/61A Undergoes Minor Structural Changes at Higher Temperature. (A) Left panel, Melting analysis of TRMT6/61A without and with tRNA. First derivatives of the fluorescence change are plotted and indicate two melting points (T_m’_ and T_m_) upon increase of the temperature. Right panel, T_m’_ is significantly lower than T_m_. ***p<0.001, two-tailed t-test; N=3 independent experiments (mean + SD). (B) Proteinase K sensitivity indicates partial unfolding of the RNA-binding TRMT6 subunit at 45°C. Right panel, the amount of protein at timepoint 0 was set as 1. **p<0.01, two-tailed t-test; N=3 independent experiments (mean + SD).

### TRMT6/61A Interacts Directly with mRNA *in Vitro*

The binding of tRNA to the TRMT6/61A complex was investigated in a direct assay *in vitro* (Figure S6A). The assay revealed an unchanged capacity of the endogenous tRNA to associate with TRMT6/61A also at higher temperature (Figure 7A). We considered the possibility that the endogenous tRNA was not a faithful substrate for the methyltransferase because it is already N1-methylated at A58. To exclude this, we next analyzed the binding of the *in vitro* synthesized, i.e., unmethylated, tRNA and again did not observe significant changes (Figure 7B).

**Figure 7.**
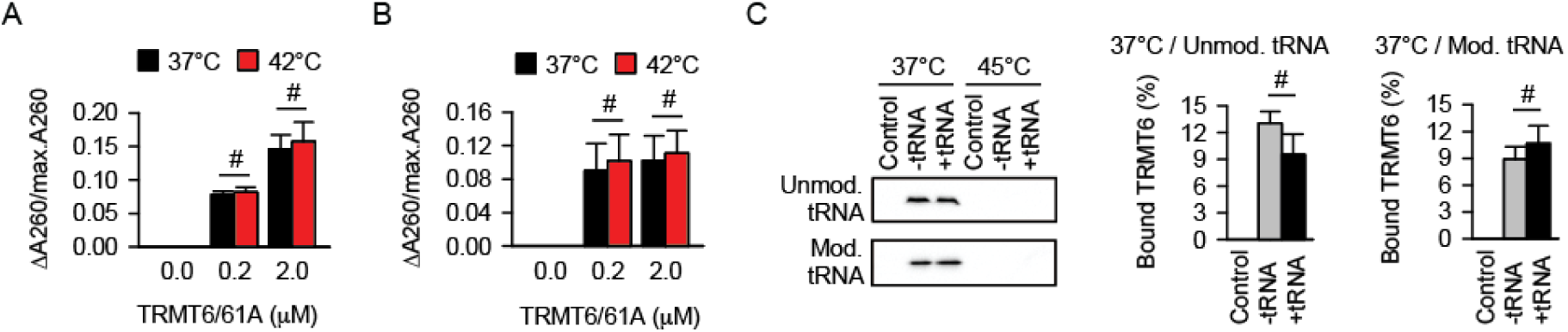
TRMT6/61A Interacts Directly with mRNA. (A and B) Comparison of TRMT6/61A binding to 1 μM endogenous tRNA (A) or 1 μM *in vitro* transcribed tRNA (B) at physiological and febrile temperature. #, not significant difference; two-tailed t-test; N=3 independent experiments (mean + SD). (C) Direct association of RNA and TRMT6/61A *in vitro*. The efficiency of TRMT6/61A pulldowns by RNA-coated beads (+/-1 μM tRNA) was estimated by western blotting. *In vitro* transcribed tRNA (Unmod. tRNA) or endogenous tRNA (Mod. tRNA) were used. Control, beads without RNA. #, not significant difference; two-tailed t-test; N=3 (mean+SD).

Finally, we succeeded to reconstitute the association of TRMT6/61A with mRNA *in vitro*. According to our estimations, the methyltransferase is found in the cytosol of HeLa cells at 1-10 nM (Figure S6B) and the tRNA concentration is known to be in low micromolar range. At these concentrations *in vitro*, very efficient direct interaction of TRMT6/61A and free RNA was detected (Figure 7C). This interaction did not depend on the presence of tRNA, regardless whether methylated or not. Contrary to the result in the lysate, increased temperature not only failed to enhance the interaction, but actually abolished it (Figure 7C).

## DISCUSSION

Inhibition of the cap-dependent translation initiation and disassembly of polysomes are two well-documented components of cellular reaction to heat shock, which are involved in stress granule (SG) formation. Our study uncovered the early step of this response in mammalian cells, namely, the capacity of mRNA - when freed from polysomes - to recruit a set of proteins involved in RNA metabolism and proteostasis. We sought to reduce sequence-specific effects by deleting 5’-UTR and 3’-UTR from both bait RNAs. Indeed, known regulation of the HSP70 mRNA operates through these regions, for example, the increased stability of the transcript ^24^ and its enhanced translation ^25^ during stress are due to the ARE in the 3′-UTR, cap-independent translation is due to IRES in the 5′-UTR ^26,27^, ribosomal shunting during heat shock operates via the 18S rRNA complementarity sequence in the 5′-UTR ^28^, transcription and translation are coupled by eEF1A1 via the 3′-UTR ^29^. It is known that HSP70 mRNA escapes TIA-positive SG ^14^. Our data indicate that the escape involves the UTRs as well because the UTR-free HSP70 RNA accumulated in granules when introduced into cells (Figure 1B).

It is not trivial to setup RNA association analyses devoid of evolutionary established sequence-specific interactions. Yet, the significant overlap of the interactomes of two different RNA sequences is reassuring that we uncovered, at least partially, the common processing machinery of free RNA released from polysomes during stress. Among interactors, there was G3BP1 known to be required for SG assembly ^30^. G3BP1 promotes phase separation in association with Caprin1 ^31^ which was also among the hits. Another indication that the interactome we report does represent an early snapshot of free RNA processing in the cytosol is the abundance of RNA helicases which would be capable of supporting the dynamic packing of long polynucleotides into SGs. At the same time, one should avoid overgeneralization of results from this kind of analysis because of the heterogeneity of SGs formed by different stressors or in different cells ^32^. Secondly, it cannot be excluded that initial stages of granulation are mechanistically similar between different types of RNA granules, for example, between SGs and P-bodies. For long, P-bodies were believed to be main sites of RNA decay, but also for storage ^33^. In this regard, the presence of the endonuclease Dicer in the free RNA interactome is especially intriguing.

Recent studies suggested that a network of proteins exists pre-formed under normal conditions and is recruited to build up granules during stress ^32,34^. If so, only a few additional components might be needed to induce granule formation. Our *in vitro* experiments using recombinant TRMT6/61A support this scenario. TRMT6/61A, one of the highest enrichments in the interactome, has been known as N1-adenine methyltrasferase of A58 in tRNAs and was recently shown to methylate mRNAs as well. Why does proteostasis stress increase the association of TRMT6/61A with mRNAs? We observed a local unfolding of the TRMT6 subunit around the temperature of the heat shock. Yet, the unfolding did not weaken its affinity towards tRNA and thus cannot explain the increase of methyltransferase binding to mRNA due to reduced competition with tRNA. We could also exclude that the local unfolding increases the affinity to mRNA since TRMT6/61 completely lost its interaction with mRNA in an *in vitro* reconstitution under higher temperature (Figure 7C). One possible explanation is that a collaborative association of several proteins with free RNA is taking place at increased temperature. A pre-existing or acutely formed network of proteins could then facilitate complex association of further proteins, such as TRMT6/61A. Thus, additional components would be needed to reconstitute the enhanced binding of some interactors to RNA upon heat shock. The nuclease sensitivity of the co-precipitation of TRMT6/61A and eIF2α supports this possibility (Figures 5A and 5B).

A number of neurodegenerative disorders are associated with repeat expansions in the disease-linked genes ^35^. As a result, faulty polypeptide products of the mutant genes misfold and aggregate sequestering a number of regulatory proteins, which interferes with diverse cellular functions ^36,37^. RNA-binding proteins constitute a significant part of coaggregating species. On the other side, repeat-containing RNAs can form disease-associated granules as well ^38^. Moreover, even bulk RNA as a biopolymer seems to modulate phase separation of prion-like proteins ^5^. Our data supports the active role of non-mutant RNA in granulation processes during proteostasis stress.

## CONCLUSIONS

The present study directly links to the recent advances in the field of RNA-protein granulation, namely, an increasing appreciation of the active role of RNAs in this process. Composition of several types of RNA-protein granules could be determined recently. The challenge now is to understand the mechanisms of their formation. Our study provides a molecular snapshot of the RNA-driven protein association early during heat shock and thus sets the ground towards this understanding. The significant overlap of the interactomes of two different RNAs argues for the general validity of the identified protein set. We hope that the data will prompt in-depth mechanistic and functional studies of individual interactors in the future.

## ASSOCIATED CONTENT

### Supporting Information

Supporting Information includes six figures (PDF) and four tables (XLSX).

**Table S1.** MaxLFQ Quantitative Data and Identifiers of mRNA (HSP70 Coding Region) Interactors.

**Table S2.** Free RNA (HSP70 Coding Region) Interactors in the Heat-Shocked Mammalian Cytosol.

**Table S3.** MaxLFQ Quantitative Data and Identifiers of Control mRNA (BRaf Coding Region) Interactors.

**Table S4.** Free Control RNA (BRaf Coding Region) Interactors in the Heat-Shocked Mammalian Cytosol.

## AUTHOR INFORMATION

### Notes

The authors declare no competing financial interest.

## ACKNOWLEDGMENTS

We thank J. Finer-Moore for the bacterial expression vector encoding human TRMT6/61A. We thank M. Joppe and F. Bourdeaux for experimental assistance. We thank M. Grininger for the access to the HPLC. We thank H. Schwalbe for critical discussion. We thank the ERC (StG-311522 to R.M.V., StG-309545 to G.G.T) and DFG (EXC115 to R.M.V) for funding. M.H. is funded by DFG CRC „Molecular Mechanisms of RNA-mediated Regulation”.

## ABBREVIATIONS

MS: mass spectrometry
RBPs: RNA-binding proteins
UTRs: untranslated regions
SGs: stress granules
GO: gene ontology
CD: circular dichroism.

## REFERENCES

(1) Brangwynne, C. P.; Tompa, P.; Pappu, R. V. Polymer Physics of Intracellular Phase Transitions. Nature Physics 2015, 11 (11), 899.

(2) Li, P.; Banjade, S.; Cheng, H.-C.; Kim, S.; Chen, B.; Guo, L.; Llaguno, M.; Hollingsworth, J. V.; King, D. S.; Banani, S. F.; et al. Phase Transitions in the Assembly of Multivalent Signalling Proteins. Nature 2012, 483 (7389), 336–340.

(3) Audas, T. E.; Audas, D. E.; Jacob, M. D.; Ho, J. J. D.; Khacho, M.; Wang, M.; Perera, J. K.; Gardiner, C.; Bennett, C. A.; Head, T.; et al. Adaptation to Stressors by Systemic Protein Amyloidogenesis. Dev. Cell 2016, 39 (2), 155–168.

(4) Langdon, E. M.; Qiu, Y.; Ghanbari Niaki, A.; McLaughlin, G. A.; Weidmann, C.; Gerbich, T. M.; Smith, J. A.; Crutchley, J. M.; Termini, C. M.; Weeks, K. M.; et al. MRNA Structure Determines Specificity of a PolyQ-Driven Phase Separation. Science 2018, 360 (6391), 922– 927.

(5) Maharana, S.; Wang, J.; Papadopoulos, D. K.; Richter, D.; Pozniakovsky, A.; Poser, I.; Bickle, M.; Rizk, S.; Guillén-Boixet, J.; Franzmann, T.; et al. RNA Buffers the Phase Separation Behavior of Prion-like RNA Binding Proteins. Science 2018, 360 (6391), 918–921.

(6) Singh, G.; Pratt, G.; Yeo, G. W.; Moore, M. J. The Clothes Make the MRNA: Past and Present Trends in MRNP Fashion. Annu. Rev. Biochem. 2015, 84, 325–354.

(7) King, O. D.; Gitler, A. D.; Shorter, J. The Tip of the Iceberg: RNA-Binding Proteins with Prion-like Domains in Neurodegenerative Disease. Brain Res. 2012, 1462, 61–80.

(8) Kato, M.; Han, T. W.; Xie, S.; Shi, K.; Du, X.; Wu, L. C.; Mirzaei, H.; Goldsmith, E. J.; Longgood, J.; Pei, J.; et al. Cell-Free Formation of RNA Granules: Low Complexity Sequence Domains Form Dynamic Fibers within Hydrogels. Cell 2012, 149 (4), 753–767.

(9) Jain, S.; Wheeler, J. R.; Walters, R. W.; Agrawal, A.; Barsic, A.; Parker, R. ATPase-Modulated Stress Granules Contain a Diverse Proteome and Substructure. Cell 2016, 164 (3), 487–498.

(10) Brandt, F.; Carlson, L.-A.; Hartl, F. U.; Baumeister, W.; Grünewald, K. The Three-Dimensional Organization of Polyribosomes in Intact Human Cells. Mol. Cell 2010, 39 (4), 560–569.

(11) Bounedjah, O.; Desforges, B.; Wu, T.-D.; Pioche-Durieu, C.; Marco, S.; Hamon, L.; Curmi, P. A.; Guerquin-Kern, J.-L.; Piétrement, O.; Pastré, D. Free MRNA in Excess upon Polysome Dissociation Is a Scaffold for Protein Multimerization to Form Stress Granules. Nucleic Acids Res. 2014, 42 (13), 8678–8691.

(12) McQuin, C.; Goodman, A.; Chernyshev, V.; Kamentsky, L.; Cimini, B. A.; Karhohs, K. W.; Doan, M.; Ding, L.; Rafelski, S. M.; Thirstrup, D.; et al. CellProfiler 3.0: Next-Generation Image Processing for Biology. PLoS Biol. 2018, 16 (7), e2005970.

(13) Silver, J. T.; Noble, E. G. Regulation of Survival Gene Hsp70. Cell Stress Chaperones 2012, 17 (1), 1–9.

(14) Kedersha, N.; Anderson, P. Stress Granules: Sites of MRNA Triage That Regulate MRNA Stability and Translatability. Biochem. Soc. Trans. 2002, 30 (Pt 6), 963–969.

(15) Gerstberger, S.; Hafner, M.; Tuschl, T. A Census of Human RNA-Binding Proteins. Nat. Rev. Genet. 2014, 15 (12), 829–845.

(16) Bolognesi, B.; Lorenzo Gotor, N.; Dhar, R.; Cirillo, D.; Baldrighi, M.; Tartaglia, G. G.; Lehner, B. A Concentration-Dependent Liquid Phase Separation Can Cause Toxicity upon Increased Protein Expression. Cell Rep 2016, 16 (1), 222–231.

(17) Holcik, M.; Sonenberg, N. Translational Control in Stress and Apoptosis. Nat. Rev. Mol. Cell Biol. 2005, 6 (4), 318–327.

(18) Oerum, S.; Dégut, C.; Barraud, P.; Tisné, C. M1A Post-Transcriptional Modification in TRNAs. Biomolecules 2017, 7 (1), pii: E20.

(19) Dominissini, D.; Nachtergaele, S.; Moshitch-Moshkovitz, S.; Peer, E.; Kol, N.; Ben-Haim, M. S.; Dai, Q.; Di Segni, A.; Salmon-Divon, M.; Clark, W. C.; et al. The Dynamic N(1)-Methyladenosine Methylome in Eukaryotic Messenger RNA. Nature 2016, 530 (7591), 441–446.

(20) Li, X.; Xiong, X.; Wang, K.; Wang, L.; Shu, X.; Ma, S.; Yi, C. Transcriptome-Wide Mapping Reveals Reversible and Dynamic N(1)-Methyladenosine Methylome. Nat. Chem. Biol. 2016, 12 (5), 311–316.

(21) Safra, M.; Sas-Chen, A.; Nir, R.; Winkler, R.; Nachshon, A.; Bar-Yaacov, D.; Erlacher, M.; Rossmanith, W.; Stern-Ginossar, N.; Schwartz, S. The M1A Landscape on Cytosolic and Mitochondrial MRNA at Single-Base Resolution. Nature 2017, 551 (7679), 251–255.

(22) Li, X.; Xiong, X.; Zhang, M.; Wang, K.; Chen, Y.; Zhou, J.; Mao, Y.; Lv, J.; Yi, D.; Chen, X.-W.; et al. Base-Resolution Mapping Reveals Distinct M1A Methylome in Nuclear- and Mitochondrial-Encoded Transcripts. Mol. Cell 2017, 68 (5), 993–1005.e9.

(23) Finer-Moore, J.; Czudnochowski, N.; O’Connell, J. D.; Wang, A. L.; Stroud, R. M. Crystal Structure of the Human TRNA m(1)A58 Methyltransferase-TRNA(3)(Lys) Complex: Refolding of Substrate TRNA Allows Access to the Methylation Target. J. Mol. Biol. 2015, 427 (24), 3862–3876.

(24) Zhao, M.; Tang, D.; Lechpammer, S.; Hoffman, A.; Asea, A.; Stevenson, M. A.; Calderwood, S. K. Double-Stranded RNA-Dependent Protein Kinase (Pkr) Is Essential for Thermotolerance, Accumulation of HSP70, and Stabilization of ARE-Containing HSP70 MRNA during Stress. J. Biol. Chem. 2002, 277 (46), 44539–44547.

(25) Moseley, P. L.; Wallen, E. S.; McCafferty, J. D.; Flanagan, S.; Kern, J. A. Heat Stress Regulates the Human 70-KDa Heat-Shock Gene through the 3’-Untranslated Region. Am. J. Physiol. 1993, 264 (6 Pt 1), L533–537.

(26) Rubtsova, M. P.; Sizova, D. V.; Dmitriev, S. E.; Ivanov, D. S.; Prassolov, V. S.; Shatsky, I. N. Distinctive Properties of the 5’-Untranslated Region of Human Hsp70 MRNA. J. Biol. Chem. 2003, 278 (25), 22350–22356.

(27) Hernández, G.; Vázquez-Pianzola, P.; Sierra, J. M.; Rivera-Pomar, R. Internal Ribosome Entry Site Drives Cap-Independent Translation of Reaper and Heat Shock Protein 70 MRNAs in Drosophila Embryos. RNA 2004, 10 (11), 1783–1797.

(28) Yueh, A.; Schneider, R. J. Translation by Ribosome Shunting on Adenovirus and Hsp70 MRNAs Facilitated by Complementarity to 18S RRNA. Genes Dev. 2000, 14 (4), 414–421.

(29) Vera, M.; Pani, B.; Griffiths, L. A.; Muchardt, C.; Abbott, C. M.; Singer, R. H.; Nudler, E. The Translation Elongation Factor EEF1A1 Couples Transcription to Translation during Heat Shock Response. Elife 2014, 3, e03164.

(30) Matsuki, H.; Takahashi, M.; Higuchi, M.; Makokha, G. N.; Oie, M.; Fujii, M. Both G3BP1 and G3BP2 Contribute to Stress Granule Formation. Genes Cells 2013, 18 (2), 135–146.

(31) Kedersha, N.; Panas, M. D.; Achorn, C. A.; Lyons, S.; Tisdale, S.; Hickman, T.; Thomas, M.; Lieberman, J.; McInerney, G. M.; Ivanov, P.; et al. G3BP-Caprin1-USP10 Complexes Mediate Stress Granule Condensation and Associate with 40S Subunits. J. Cell Biol. 2016, 212 (7), 845–860.

(32) Markmiller, S.; Soltanieh, S.; Server, K. L.; Mak, R.; Jin, W.; Fang, M. Y.; Luo, E.-C.; Krach, F.; Yang, D.; Sen, A.; et al. Context-Dependent and Disease-Specific Diversity in Protein Interactions within Stress Granules. Cell 2018, 172 (3), 590–604.e13.

(33) Hubstenberger, A.; Courel, M.; Bénard, M.; Souquere, S.; Ernoult-Lange, M.; Chouaib, R.; Yi, Z.; Morlot, J.-B.; Munier, A.; Fradet, M.; et al. P-Body Purification Reveals the Condensation of Repressed MRNA Regulons. Mol. Cell 2017, 68 (1), 144–157.e5.

(34) Youn, J.-Y.; Dunham, W. H.; Hong, S. J.; Knight, J. D. R.; Bashkurov, M.; Chen, G. I.; Bagci, H.; Rathod, B.; MacLeod, G.; Eng, S. W. M.; et al. High-Density Proximity Mapping Reveals the Subcellular Organization of MRNA-Associated Granules and Bodies. Mol. Cell 2018, 69 (3), 517–532.e11.

(35) La Spada, A. R.; Taylor, J. P. Repeat Expansion Disease: Progress and Puzzles in Disease Pathogenesis. Nat. Rev. Genet. 2010, 11 (4), 247–258.

(36) Schaffar, G.; Breuer, P.; Boteva, R.; Behrends, C.; Tzvetkov, N.; Strippel, N.; Sakahira, H.; Siegers, K.; Hayer-Hartl, M.; Hartl, F. U. Cellular Toxicity of Polyglutamine Expansion Proteins: Mechanism of Transcription Factor Deactivation. Mol. Cell 2004, 15 (1), 95–105.

(37) Kim, Y. E.; Hosp, F.; Frottin, F.; Ge, H.; Mann, M.; Hayer-Hartl, M.; Hartl, F. U. Soluble Oligomers of PolyQ-Expanded Huntingtin Target a Multiplicity of Key Cellular Factors. Mol. Cell 2016, 63 (6), 951–964.

(38) Jain, A.; Vale, R. D. RNA Phase Transitions in Repeat Expansion Disorders. Nature 2017, 546 (7657), 243–247.

