## Supplemental Information for "Active role of free RNA during the early phase of proteostasis stress"

- **Figure S1.** RNA and Proteins Form Granules in HeLa Cells.
- **Figure S2.** Identification of Free RNA Interactors in Heat-Shocked Cytosol.
- **Figure S3.** Free RNA Interactors Differ from the Conventional RBPs.
- **Figure S4.** Translation Machinery is Affected by Heat Shock.
- **Figure S5.** *In Vitro* Characterization of Recombinant TRMT6/61A.
- **Figure S6.** RNA Binding to Recombinant TRMT6/61A Can Be Detected *In Vitro*.

- **Table S1.** MaxLFQ Quantitative Data and Identifiers of mRNA (HSP70 Coding Region) Interactors.
- **Table S2.** Free RNA (HSP70 Coding Region) Interactors in the Heat-Shocked Mammalian Cytosol.
- **Table S3.** MaxLFQ Quantitative Data and Identifiers of Control mRNA (BRaf Coding Region) Interactors.
- **Table S4.** Free Control RNA (BRaf Coding Region) Interactors in the Heat-Shocked Mammalian Cytosol.

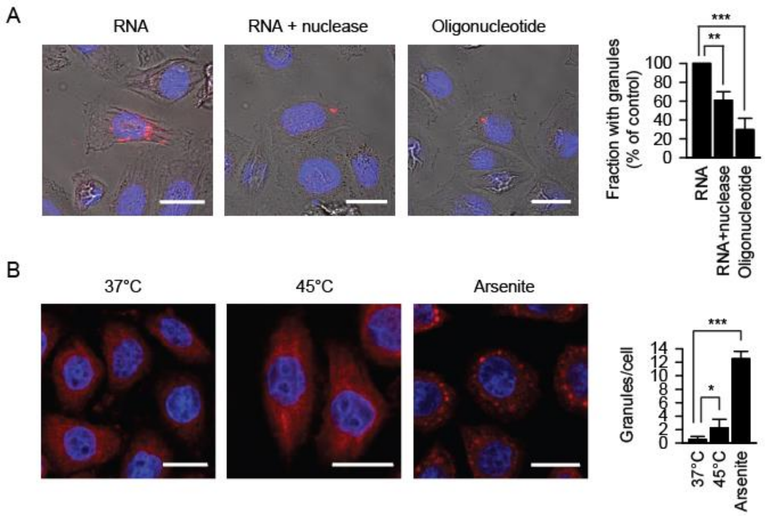


**Figure S1. RNA and Proteins Form Granules in HeLa Cells.**

(A) HeLa cells 6 h after electroporation with Cy5 (red) 5'-labelled RNA (RNA) or 55mer oligomer. One part of the RNA transfectants were treated with nuclease before imaging (RNA + nuclease). DAPI staining (blue), nuclei. Scale bar 20 μm. Merged images are shown. **p<0.01, ***p<0.001, two-tailed t-test; N=3 independent experiments (mean + SD).

(B) Immunofluorescence staining of eIF4E (red) to detect stress granule formation upon 45°C for 1 hr or 1 mM arsenite treatment for 30 min. DAPI staining (blue), nuclei. Scale bar 20 μm. *p<0.05, ***p<0.001, one-tailed t-test; N=3 independent experiments (mean + SD).

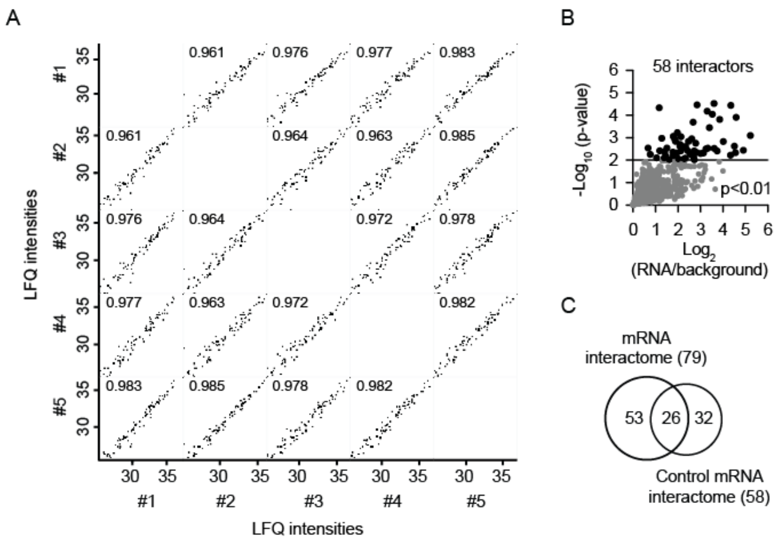


**Figure S2. Identification of Free RNA Interactors in Heat-Shocked Cytosol.**

(A) Correlation of LFQ values of quantified proteins in five independent pulldowns. The values in individual dot plots represent the squared Pearson correlation coefficients.

(B) Volcano plot of quantified proteins plotted according to their enrichment on control RNA over background with the statistical significance of the respective ratios plotted on the y-axis. A line at p-value used to define the 58 interactors (black symbols) is indicated. N=3 independent experiments.

(C) Highly significant overlap ($p<4.7 \times 10^{-63}$) between interactor sets of two RNAs in heat shocked HeLa lysates.

A

| Category value | Enrichment | P value |
|---|---|---|
| Nucleic acid binding | 2.8 | $7.95 \times 10^{-11}$ |
| RNA binding | 3.5 | $1.55 \times 10^{-9}$ |
| Nucleolus | 3.4 | $1.24 \times 10^{-7}$ |
| Non-membrane-bound organelle | 2.1 | $5.73 \times 10^{-7}$ |
| Nuclear part | 1.8 | $1.09 \times 10^{-6}$ |

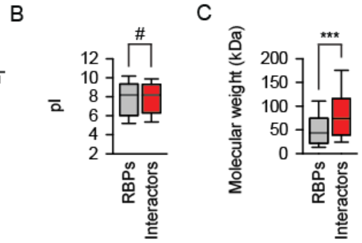


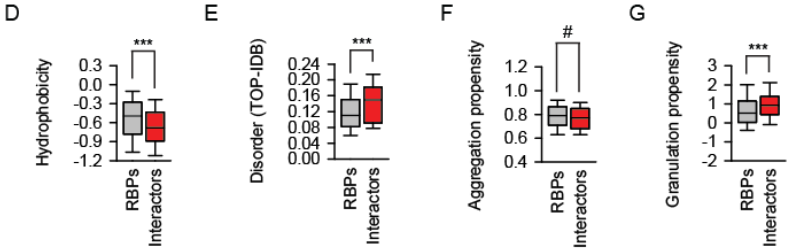


**Figure S3. Free RNA Interactors Differ from the Conventional RBPs.**

(A) GO categories enriched in the RNA interactome with the statistical significance of $p<0.001$.

(B-G) Comparison of the identified RNA interactors with human RBPs. Distribution of isoelectric points (B), molecular weights (C), Kyte-Doolittle hydrophobicity scores (D), predicted disorder (E), aggregation propensity (F) and granulation propensity (G) scores of the indicated sets of proteins. ***$p<0.001$, Mann-Whitney test. #, not significant difference.

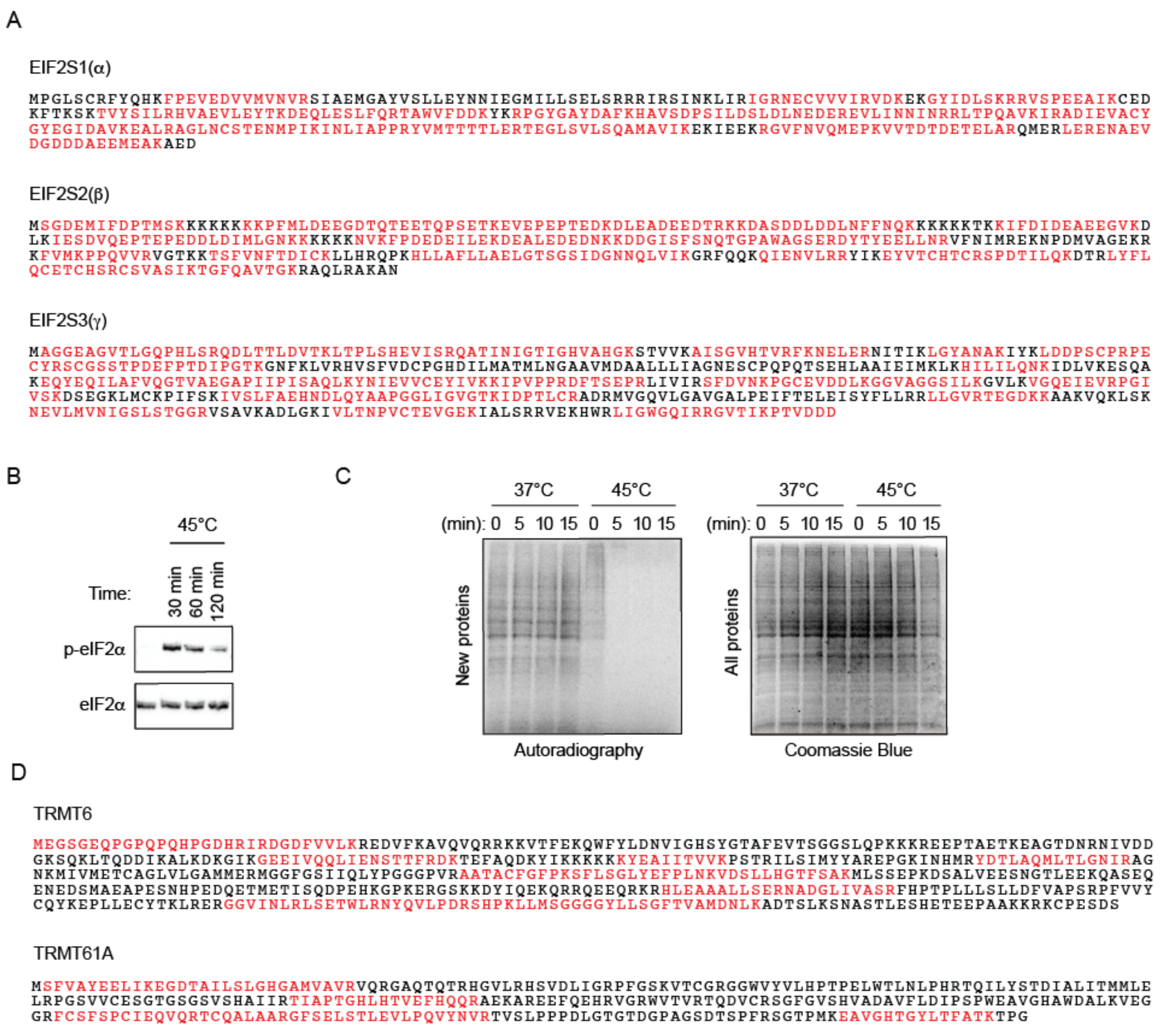


**Figure S4. Translation Machinery is Affected by Heat Shock.**

(A) Sequence coverage (red) during mass spectrometry analysis of the indicated subunits of the eIF2 complex.

(B) eIF2α serine 51 phosphorylation (p-eIF2α) in HeLa cells during heat shock detected in western blotting with phosphosite-specific antibody. In parallel, the total amount of eIF2α was analyzed. One representative out of three experiments is shown.

(C) Analysis of protein synthesis during heat shock in HeLa cells by means of radioactive labeling of newly translated polypeptides, SDS-PAGE separation and autoradiography (left panel). Right panel, Coomassie blue staining of SDS-PAGE gel to detect total amount of proteins. The same gel was used for both, autoradiography and dye staining. One representative out of three experiments is shown.

(D) Sequence coverage (red) during mass spectrometry analysis of the indicated subunits of the methyltransferase TRMT6/61A.

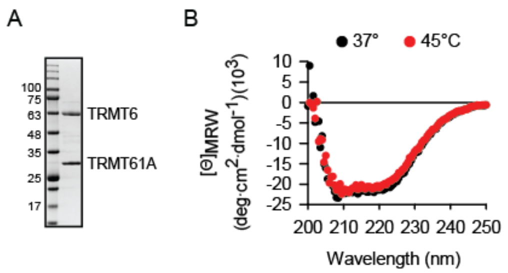


**Figure S5. *In Vitro* Characterization of Recombinant TRMT6/61A.**

(A) Coomassie Blue staining of SDS-PAGE gel to analyze the purity of recombinant methyltransferase TRMT6/61A.

(B) CD spectroscopy of 5 μM TRMT6/61A solution to analyze the secondary structure of the complex at 37°C (black) and 45°C (red). One representative out of three independent experiments is shown.

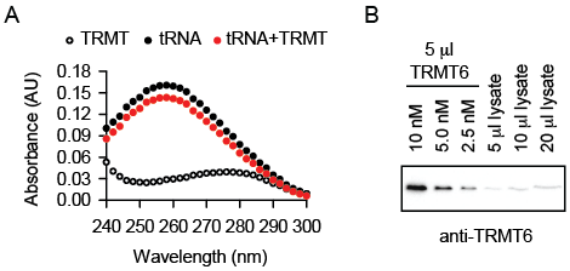


**Figure S6. RNA Binding to Recombinant TRMT6/61A Can Be Detected *In Vitro*.**

(A) Absorbance scan of 2 µM TRMT6/61A (white), 1 µM tRNA (black) and the mixture of 1 µM tRNA and 2 µM TRMT6/61A (red) at 37°C. One representative out of three independent experiments is shown.

(B) Western blotting to determine TRMT6 concentration in HeLa lysates. 5 µl of recombinant TRMT6 at decreasing concentration were loaded on the same gel for comparison. Please note that lysate represents ca. 10x dilution of the cytosol as estimated by the volume of cell pellet. One representative out of three independent experiments is shown.
